# Tumor Spheroid Uptake of Fluorescent Nanodiamonds is limited by Mass Density: a 4D Light-Sheet Assay

**DOI:** 10.1101/2024.11.15.623765

**Authors:** Maria Niora, Martin Dufva, Liselotte Jauffred, Kirstine Berg-Sørensen

## Abstract

Fluorescent nanodiamonds (FNDs) with nitrogen-vacancy centers are promising candidates for long-term biolabeling and biosensing applications due to their biocompatibility and unique optomagnetic properties. The employment of nanomaterials in cancer therapy and diagnostics, requires a deep understanding of how nanoparticles (NPs) interact with the three-dimensional (3D) tumor environment. We developed a novel approach, the “Tumor-in-a-Tube” platform, using 4D light-sheet microscopy to explore the spatio-temporal dynamics of FNDs with 3D tumor spheroids. By monitoring the real-time NP sedimentation, spheroid penetration, and cellular uptake of FNDs and polystyrene nanoparticles (PNPs), we marked the impact of NP mass density on their spheroid interaction. Unlike PNPs, higher-density FNDs underwent rapid sedimentation, which minimized their effective concentration and hindered the FND – spheroid interactions. This results in constrained intratumoral accumulation, and size-independent uptake and penetration. Longer FND effective-exposure-time promotes size-dependent cell uptake, verified by FND treatment on 2D monolayers. Nonetheless, FNDs exhibited good biocompatibility and long-term spheroid labeling, allowing for cell isolation from different spheroid layers. Our results suggest the need for NP effective-exposure-time calibration in comparative NP assays, in 3D static models. Overall, our platform provides a valuable tool for bridging the gap between 2D and 3D static models in NP assessment, drug delivery, toxicology profiling and translational research.

## Introduction

Nanotechnology is a rapidly evolving tool in the fields of cancer therapy, exploring nanomaterials with potential in enhancing therapeutic outcomes and diagnostic imaging. Fluorescent nanodiamonds (FNDs) with nitrogen-vacancy (NV) centers hold great promise as safe, stable and efficient agents. FNDs are carbon particles in the nanoscale with native fluorescence, which is spin-dependent and qualifies them in various biomedical applications. They are increasingly explored *in vitro* and they have been featured as long-term cell labels ^1–4^, sensors of biological processes ^5–9^, drug carriers ^10–12^ and biomarkers ^3,13,14^.

A major challenge in NP-based cancer therapy is achieving effective delivery of drug and diagnostic agents to tumors. Three dimensional (3D) *in vitro* tumor spheroid models are widely approved to evaluate nanocarrier efficiency. Compared to traditional 2D cell cultures, multicellular spheroids better mimic *in vivo* tumor complexity in architecture, cellular heterogeneity and gradients of nutrient, oxygen, and NP penetration ^15,16^. FNDs offer superior optical properties for long-term spheroid labeling, compared to conventional fluorophore-conjugated NPs, due to the non-blinking, non-bleaching, stable FND fluorescence from nitrogen-vacancy (NV) centers embedded in their crystal lattice. Besides their fluorescence properties, FNDs have well-documented biocompatibility^3,17–21^ and can be used as intracellular nanoscale magnetic sensors ^22–26^, offering diagnostic and theranostic promise^27^. Yet, within 3D tumor models, FNDs remain underexplored ^4,28^. Treatment of tumor spheroids with small drug-loaded nanodiamonds is reported to reduce drug resistance in the long-run^29^, through sustained drug release^30,31^, however sometimes together with limited spheroid penetration to the spheroid rim^28^, which is dependent to both nanoparticle properties and the spheroid model.

The interaction of NPs with the tumor cells, especially within 3D environments, remains complex and poorly understood. 3D spheroids are a good model for studying the distribution profile of the NPs within a tumor and explore the NP properties, such as the well-documented size, shape, surface charge and applied dose, that affect their efficiency and potential toxicity^32,33^. However, static 3D models cannot visualize the dynamic NP – spheroid interactions, which affect NP penetration, uptake, and accumulation. Real-time platforms, like organ-on-a-chip systems, enable dynamic tracking of these interactions in controlled microenvironments, surpassing static 2D and 3D assessment methods.

In this study, we developed a platform of similar concept, “Tumor-in-a-Tube”, by adopting Airy-beam light-sheet microscopy for real-time imaging of FND penetration into 3D tumor spheroids. To our knowledge, this is the first study on the spatio-temporal dynamics of such nanodiamonds in tumor spheroids. Our approach unveils the early NP – spheroid interaction and the often overlooked impact of mass density on the NP – cell interactions, which may result in contradicting results with 2D cultures, particularly in toxicology profiling^34^. We investigate the distribution profile, retention and uptake of commercial carbocylated FNDs of two different diameters, i.e., 70nm and 120nm, which is an optimal size range in bio-sensing applications, based on ongoing research in our group. FNDs of both sizes effectively penetrate the first tumor surface layers showing potential in screening assays, however with significantly lower accumulation compared to polystyrene nanoparticles (PNPs).

Our platform reveals that high NP mass density weakens spheroid uptake, due to the rapid sedimentation that reduces their effective availability for tumor interactions. These insights are valuable in NP design for improved therapeutic strategies and bring attention to how NP mass density differentially affects NP – cell interactions on 2D vs static 3D models. With protocol modifications to compensate for their short effective-exposure-time in 3D static models, FNDs can serve as inert label-probes for long-term tumor monitoring, and could be implemented as theranostic agents and in 3D biosensing experiments. Finally, our methods can easily be adapted to *in vitro* drug screening, to facilitate NP performance controls and dose optimization.

## Results and discussion

### The “Tumor-in-a-Tube” Platform

A schematic illustration of the tumor-in-a-tube platform and the investigated dynamic concepts are depicted in Figure 1. A tube embedding a tumor spheroid is mounted on a light-sheet microscope for live cell time-lapses. Nanoparticles (NPs) are introduced to the spheroid and two channels of fluorescence are recorded; one channel detects the NPs and the other localizes the spheroid within the tube to facilitate image analysis. Upon reaching the spheroid, the NPs may either interact with the spheroid cells on the surface layers or pass through the intercellular spaces and reach deeper spheroid layers (Fig.1B). At the single cell level, the NPs can adhere to the plasma membrane, followed by potential agglomeration or cellular uptake via physiological pathways (Fig.1C). Last, the free NPs in the surrounding medium may undergo sedimentation during the experiment (Fig.1D).

**Figure 1.**
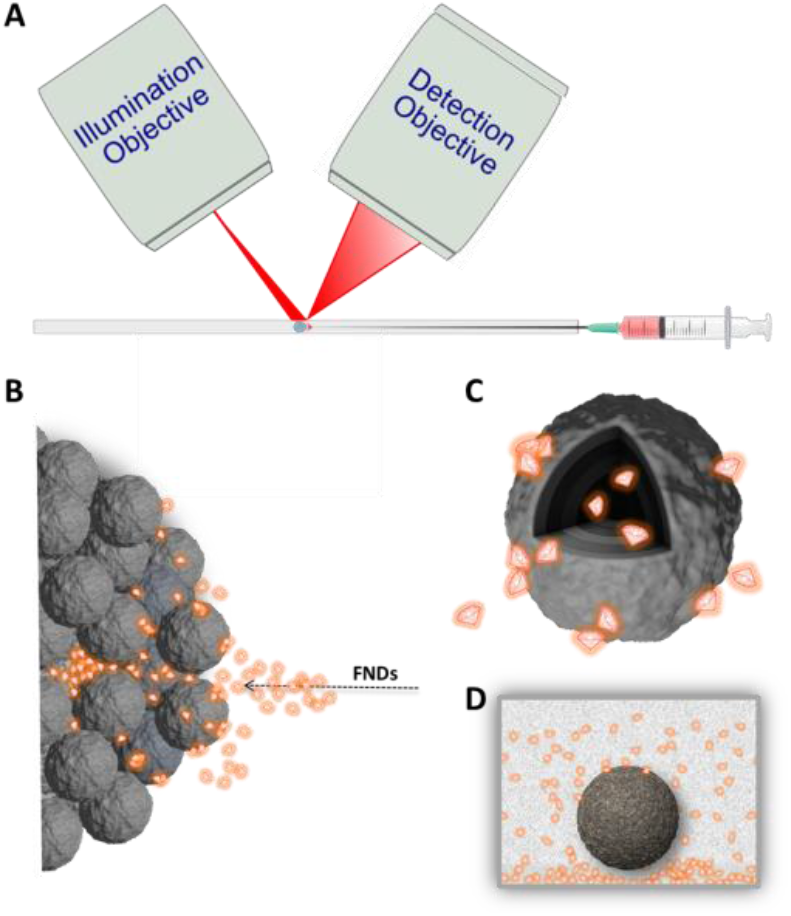
“Tumor-in-a-Tube” platform. A) A tumor spheroid is positioned inside a tube mounted on a light-sheet microscope. Fluorescent nanodiamonds (FNDs) are injected and the dynamic interactions between the FNDs and the spheroids are captured through time-lapse imaging. FNDs can B) surround and penetrate the spheroid or interact with the spheroid cells by C) adhering to the plasma membrane, where they may accumulate or get internalized by physiological uptake mechanisms. D) FNDs in suspension and under sedimentation at the bottom of the tube.

### Temporal Accumulation and Uptake of FNDs and PNPs in U87 Spheroids

The tumor spheroids were cultivated from U87 glioblastoma multiforme cells following the methodology described before^35^. In accordance with previous works, after three days of culture in ultra-low attachment wells (5,000 cells / mL), viable spheroids with a defined border and without a necrotic core are formed^35–38^ (Fig.S1). These spheroids reached an average radius of 123 ± 9 μm and a roundness of 0.8 ± 0.04 (SD, n = 14, CV ≤ 1) by means of circularity, while individual U87 cells in the spheroid arrangement measured approximately 19 ± 4 μm in diameter (SD, n = 50).

We compared the spheroid uptake of commercial fluorescent nanodiamonds (FNDs) to widely studied commercial polystyrene nanoparticles (PNPs) ^39–42^. Both FNDs and PNPs had carboxylated surface and ζ-potential around -40 mV^14,43^ and -30 mV^35,44^, respectively, as per manufacturer data and literature. They differ in shape with spherical PNPs^42^ versus flake-like shaped FNDs^45,46^ and in nominal size: 20 nm (PNP20), 70 nm (FND70), 100nm (PNP100) and 120 nm (FND120). These NPs reflect the preferred material and size range for cancer research^47,48^ and the FND size range is additionally suitable for intracellular magnetometry. We also tested 30 nm FNDs, but we could not resolve them with our methods.

The temporal accumulation of FNDs and PNPs in U87 tumor spheroids is shown in Figure 2. The maximized projections of NP signal enable visualization of both NPs at the spheroids and NPs in suspension. Five minutes after administration, PNP20, FND120 and FND70 are observed at the spheroids while all NPs are monitored at the background (Fig.2A). Two hours later, all NP types accumulated in the spheroids with the greatest fluorescence exhibited by PNP20. Owing to their lower number of nitrogen-vacancy centers, FND70 fluoresce significantly lower than FND120, at low particle concentrations. No FNDs are visible in suspension, due to severe sedimentation, contrary to the PNPs (Fig.2B). At the single cell level, PNPs displayed more uniform distribution than FNDs (Fig.2C). PNPs were detected both at the plasma membranes and intracellularly, but subcellular localization of the dimmer FNDs was challenging.

**Figure 2.**
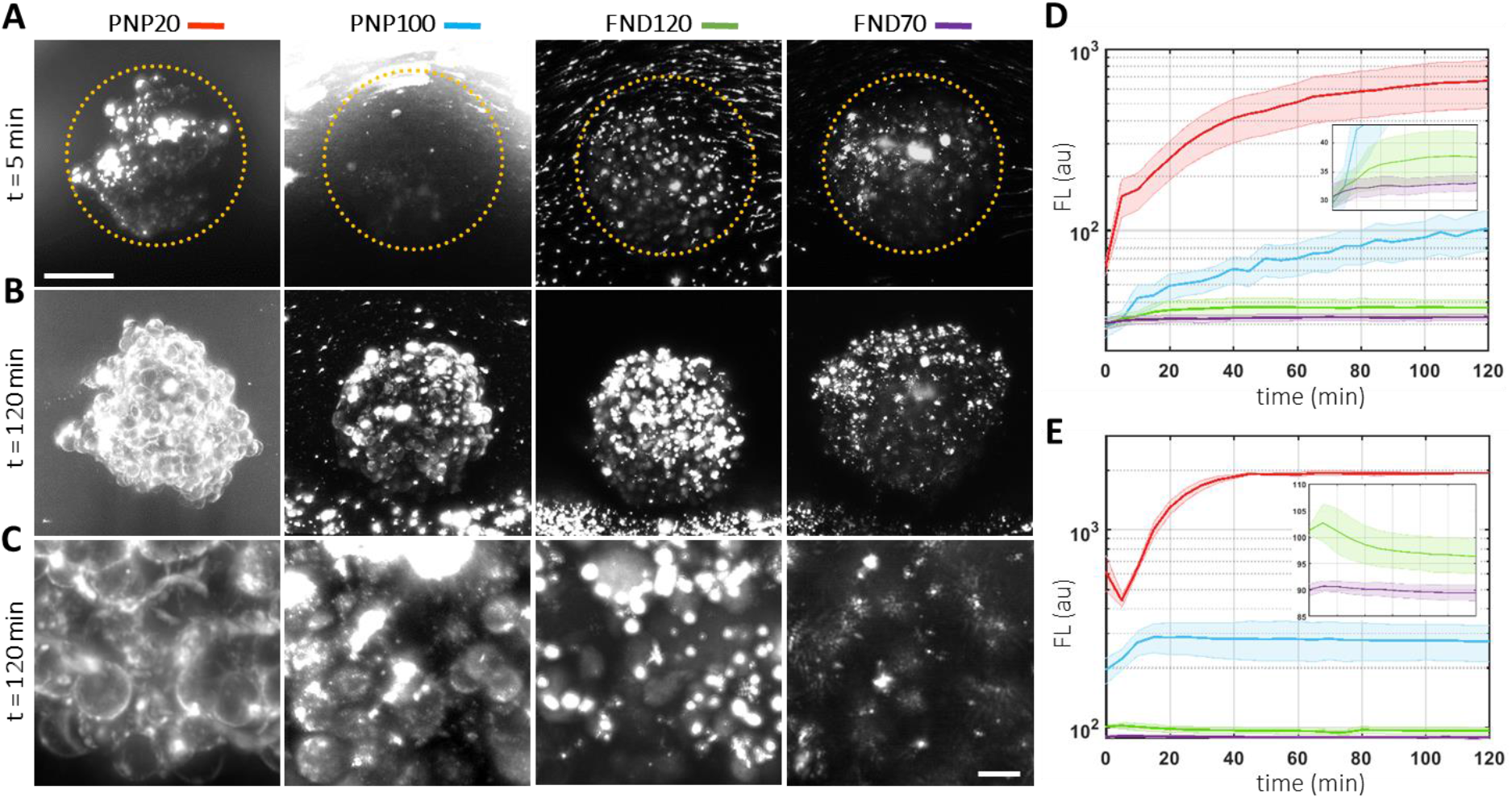
Nanoparticle accumulation and uptake in tumor spheroids. Maximized intensity projections at A) t= 5 min versus B) t= 120 min after application of polystyrene nanoparticles with diameter 20 nm (PNP20, red) or 100 nm (PNP100, blue) or fluorescent nanodiamonds with diameter 120 nm (FND120, green) or 70 nm (FND70, purple). The dashed circles indicate the spheroid position. Scale bar: 100 μm. C) Zoom-in at the surface layers. Scale bar: 15 μm. D) Integrated densities of the fluorescence signal within the tumor spheroids and E) mean background fluorescence surrounding the spheroid surface, over time. The insets zoom in to the FNDs during the first 60 min (SD, n= 3).

To illustrate the timescale of the NP-spheroid interaction, we measured NP signal intensities (FL) in the spheroids and at the background over time. The integrated densities elucidate the varying NP accumulation within the tumors with *FL*_*PNP20*_ >> *FL*_*PNP100*_ >> *FL*_*FND120*_ > *FL*_*FND70*_ (Fig.2D). PNP20 outperformed all NPs, while FND70 demonstrated the lowest signal, even compared to their larger counterparts FND120, although without statistical significance (p = 0.855, one-way ANOVA). The background fluorescence intensity (*BFI*) matches this trend, with *BFI*_*PNP20*_ >> *BFI*_*PNP100*_ >> *BFI*_*FND120*_ > *BFI*_*FND70*_ (P_FND70−FND120_ < 0.01, one-way ANOVA) (Fig.2E). *BFI*_*PNP*_ reached a plateau that is consistent to the observation of PNPs in suspension throughout the experiment, unlike *BFI*_*FND*_ that drops within 10 min post application. The contrast in NP availability is reflected in the NP – spheroid interactions and is attributed to the mass density of the nanoparticles that significantly affects the NP effective-concentration (Figure S2).

### The Effect of FND Mass Density on the FND – Spheroid interaction

To understand the impact of NP mass density (*ρ*) on their availability to the spheroids, we estimated the effect of gravity on their travelling distance (*d*), based on Stoke’s law that describes the sedimentation of spherical particles in a fluid (Equation 1). A simple estimate of *d* due to gravitational settling over a given timespan is derived from the balance between frictional and gravitational forces, determining the settling velocity (*v*) of a single NP. At low Reynolds number dynamics inertia is negligible. Using this approach, *d* within a given timespan scales with the difference in density between the NP and the surrounding medium, (*ρ*_*NP*_ − *ρ*_*medium*_), and with the NP radius squared (*r*^*2*^). Given the similar densities of PNPs and the surrounding medium (*ρ*_*PNP*_ ≈ *ρ*_*medium*_), with *ρ*_*PNP*_ = 1.05 g/cm^3^, versus the relatively higher *ρ*_*FND*_ = 3.5 g/cm^3^, then after 60 min, *d*_*PNP2*0_ is roughly just 0.04 μm, whereas *d*_*FND*1*2*0_ reaches 72 μm. This means that FND120 sediment about 1800 times faster, thereby temporally limiting their interactions with the spheroids. The estimated distances for PNP100 and for FND70 are 1 μm and 24 μm, respectively. The effect is greater with NP dispersion and susceptibility to aggregation, while other factors such as the NP shape, the Brownian motion, and the interparticle interactions need to be considered for more precise calculations^49^. Single particle analysis on the background revealed that FND70 and FND120 peak at t = 5 min post administration and decline thereafter, with the number of detected particles in suspension dropping below 5%, relative to their peak, by t=35 min and t=25 min, respectively.

### 4D Penetration and Retention of FNDs and PNPs in U87 Spheroids

To explore the spatio-temporal penetration of the different NPs within the tumor spheroids, we employed a 3D radial distribution analysis, expanding the 2D radial analysis reported earlier ^35^. Here, the third dimension proves critical to compensate for non-uniform NP distribution at the early time-points. We confined our analysis to the quarter of the spheroid that is both closest to the objectives^35^ and to the needle that introduces the NPs in the axial-direction.

Figure 3A collectively illustrates representative optical cross-sections of the spheroids after 2 h of exposure to FND or PNP. To mitigate the variability in NP brightness, fluorescence was visually normalized with a heatmap color scheme, by applying a fire LUT with Fiji software (Fig.3B). PNP20 exhibit the most gradual decrease across *r*_*100*_ = 100um from the spheroid periphery, compared to PNP100, FND70 and FND120. FND70 and FND120 demonstrate a more restricted penetration profile than PNPs, while FND70 display the least uniform peritumoral distribution. In the background, FND70 and FND120 are non-visible, in contrast to PNP20 and PNP100, which however showcase different dispersity.

**Figure 3.**
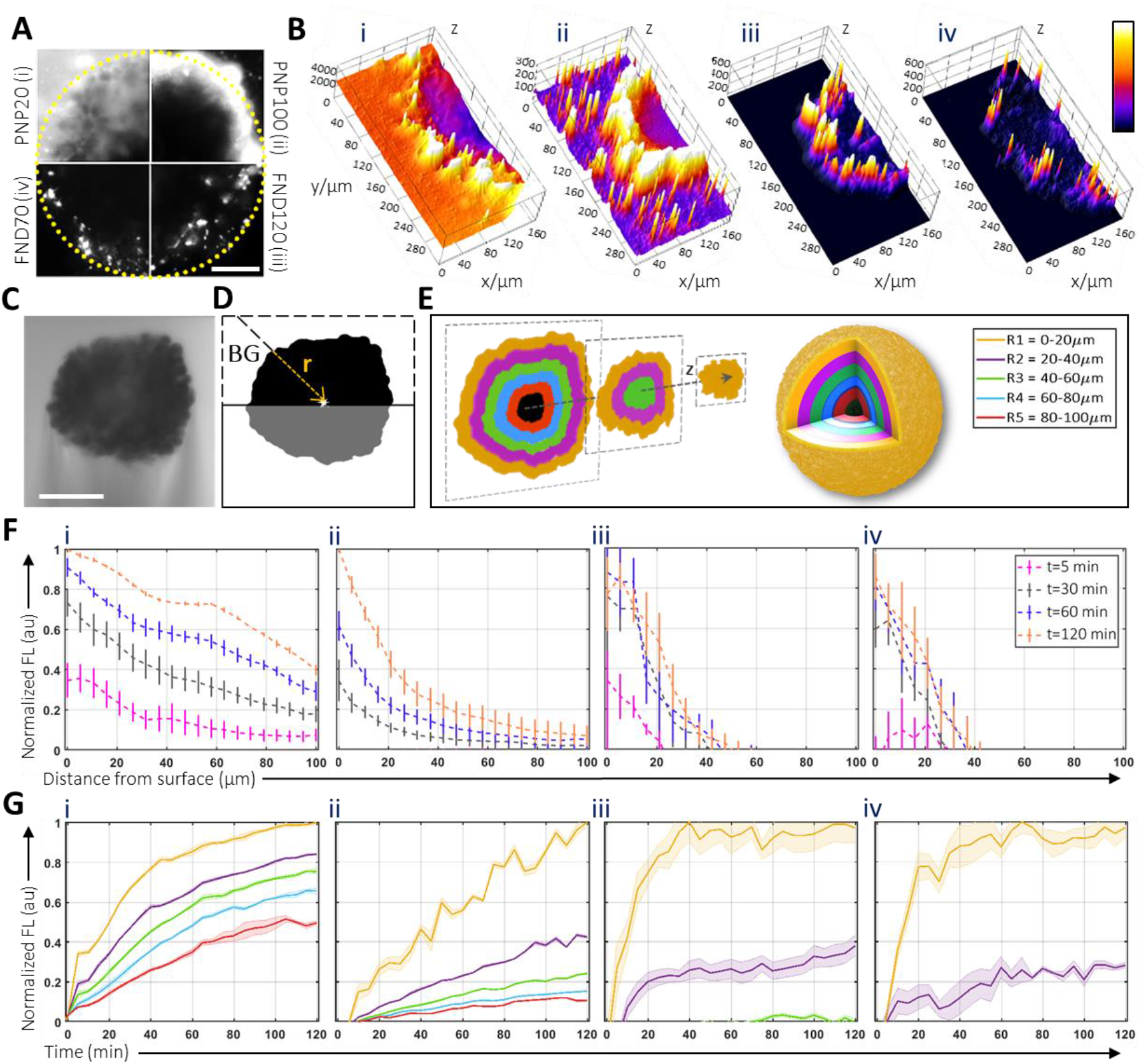
Spatio-temporal radial distribution profiles of fluorescent nanodiamonds (FNDs) and polystyrene nanoparticles (PNPs) in U87 spheroids. The nanoparticles were administered at t = 0 min and imaged every 5 min, for 2 h, with an airy-beam light sheet microscope. A) Mid-plane optical cross sections of spheroids treated for 2 h with PNPs of diameters i) 20 nm (PNP20), ii) 100 nm (PNP100) or FNDs of diameters ii) 120 nm (FND120), iv) 70 nm (FND70). Scale bar: 50 μm. B) The respective surface plots of the half side of the spheroids subjected to analysis. C) Un-stained spheroid in the fluorescent background used for generation of D) a binary image used to restrain the radial analysis to the spheroid boundaries, for radius *r*_*100*_ = 100 μm towards the core within an azimuth angle of 180°. Scale bar: 100 μm. This was repeated across the z-axis of the spheroids for a 3D perspective followed by a E) 3D shell-based assessment of the integrated signal within five regions of 20 μm intervals. 4D radial analysis across F) *r*_*100*_ in 3D at four different timepoints and G) the five 3D shells during 2 h (SD, n= 3).

For 4D radial analysis on live spheroids, precise spheroid localization within the field of view is essential. To achieve this, we opted for a cell impermeable dye that provides the necessary contrast between a brightly fluorescent background and the unstained spheroid, thereby avoiding inducing phototoxicity with a cell dye alternative (Fig.3C). Exploiting this channel post processing, we generate a time-series of 3D binary masks that help outline the spheroid periphery (Fig.3D, E).

Quantitatively, from the spheroid edge towards the core, we calculated the average fluorescence intensity across 100 μm in 3D (Fig.3F) and further assessed the integrated densities in 20 μm intervals (Fig.3G, E). Our results are normalized per NP type for easier comparison of the penetration efficacy, but the raw data indicated the same fluorescence intensity trend as in Fig. 2D. To decouple FND signal from cell autofluorescence, all data were normalized by subtracting the 4D radial intensity profiles of control spheroids subjected to timelapse imaging without NPs. FND70 and FND120 express similar spatio-temporal distribution patterns, with signal confined to roughly the first 40 μm and 50 μm from the periphery, respectively (regions R1 and R2 in Fig. 3E). Primarily as a consequence of the FND sedimentation, both sizes reached a plateau in R1 by around 60 min and 40 min respectively, signifying good retention. In contrast, PNPs present both enhanced and size-dependent penetration, with the steepest profile for PNP100. PNP20 are significantly the most efficient and rapid on reaching deeper spheroid layers (R4 and R5 in Fig3E), indicating both cell uptake and free PNP20 available for interstitial transport. As a result from the abundance of PNPs in the background surrounding the spheroids, PNP100 display sustained uptake at the outer spheroid layers (R1), but without as much convection to deeper regions.

Our results on PNPs align with similar studies reporting size-dependent penetration profiles for anionic, carboxylated PNPs^41^. PNP100 effectively reach deep spheroid layers but with reduced permeability across the spheroids is also supported^42,50^. Similarly, the size-dependency to rapid penetration into tumor spheroids and enhanced cellular uptake are documented with other types of NPs by various reports, where size was the main variable^51–53^ and including real-time platforms^37,54^. On the contrary, FNDs did not follow a size-dependent distribution likely due to their uneven intrinsic fluorescence, which for FND70 is 6-fold lower than FND120. This factor, combined with the competition from the background noise, imply that higher FND70 agglomeration is necessary before achieving detectable levels, as also noted in other works^55^. Cell autofluorescence in the spectral region of 500-600 nm is likely due to flavoproteins and is an acknowledged limitation when measuring weak fluorescent probes like the FNDs^56^. To overcome this limitation and obtain enhanced uniform FND accumulation in 3D spheroids, we suggest sequential dosing, frequent manual stirring or a rocking stage during FND application on static 3D models. We anticipate that this protocol adjustment would improve cell labeling and would likely lead to increased detected penetration. Other more elaborate methods are also reported^57^. However, for biosensing applications with magnetometry by single-particle interrogation, distinct FND particles are preferred, suggesting that even short exposure times, as brief as 30 minutes, may suffice. Nonetheless, NP penetration is also affected by the spheroid properties, such as cell stiffness and spheroid density, which are determined by the cell origin of the spheroid itself ^58^.

### Uptake and Biocompatibility of FNDs in U87 spheroids

To examine the cellular uptake of FNDs across the tumor layers, we conducted single cell analysis using FACS on dissociated spheroids following 2 h incubation with FND70 or FND120, co-labeled with a nuclear dye (Fig.4A). Spheroid dissociation strips off unbound NPs in the interstitial space and loosely bound NPs from the plasma membrane of the cells, revealing cell uptake. The nuclear labeling displayed a diffusion gradient that was used to define three spheroid regions based on the dye intensity (Fig.4B), consistent with previous works ^29,43^. FND70 and FND120 displayed similar spheroid labeling (p = 0.782) which was significantly confined to the spheroid surface layers (p < 0.0001).

**Figure 4.**
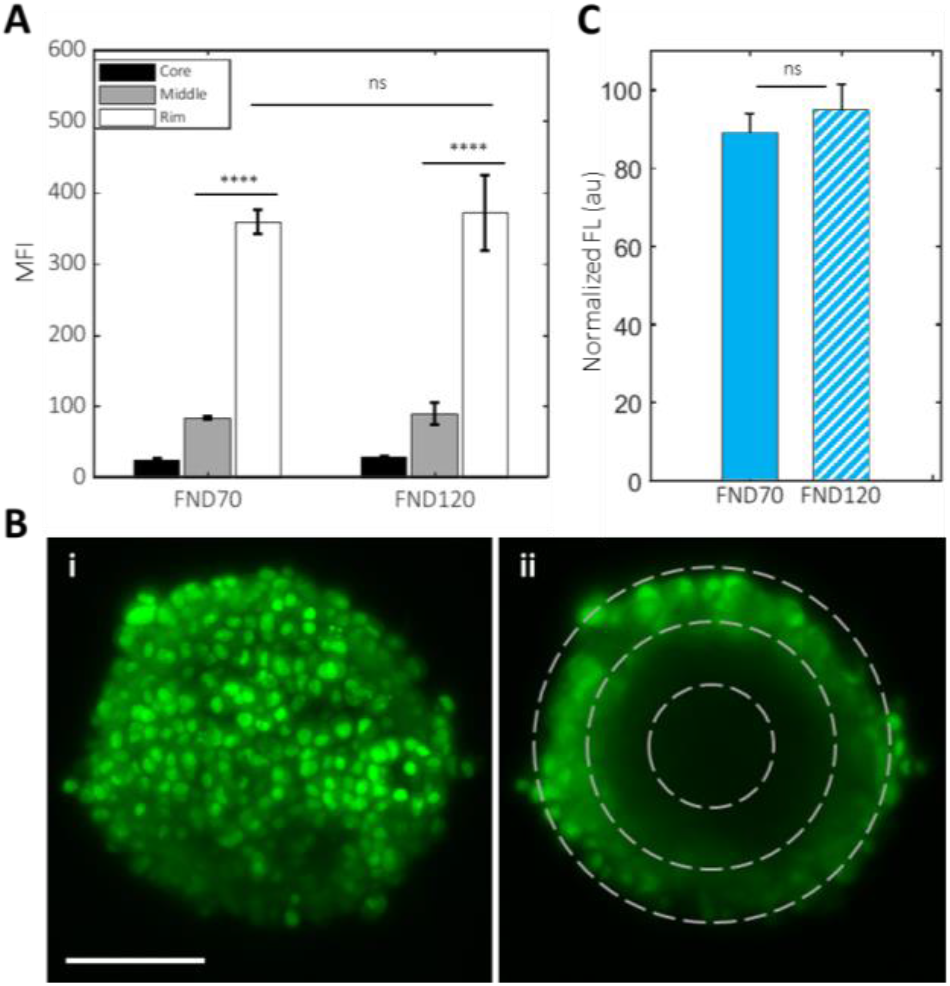
Uptake of fluorescent nanodiamonds (FNDs) in U87 tumor spheroids, after 2 h exposure. Spheroids were co-stained for nuclei and then dissociated. A) Flow cytometric analysis of FND signal in different spheroid regions, which were defined by nuclear fluorescence intensity. B) Representative nuclear labeling of a spheroid by i) maximized intensity projection and ii) at a single optical slice mid-plane. The dashed rings are indicative of the three spheroid regions used in the analysis. C) FND biocompatibility for the two FND sizes (70 nm and 120 nm) in spheroids co-stained for viability. Data are normalized by the mean value from spheroids not treated with FNDs (SD, n=3). Statistical significance: one-way ANOVA, Student’s t-test, a = 0.05, ****p < 0.0001. Scale bar: 100 μm.

For evaluation of FND biocompatibility, spheroids were additionally stained with calcein viability dye before harvesting. Control spheroids were not treated with FNDs. Treatment with FNDs did not affect the U87 spheroid viability, although FND70 showed minor viability decrease without statistical significance (p = 0.256), indicating the overall good FND biocompatibility.

### Uptake of FNDs in U87 cell monolayers

The confined accumulation of FNDs at the spheroid rim, with comparable cellular uptake for both FND sizes, prompted us to explore FND uptake on U87 cell monolayers. Confocal microscopy imaging revealed that U87 cells readily scavenge the surrounding FND120 in the course of 2 h and that FND120 exhibit efficient cell internalization by 30 min exposure (Fig.5A). The close up in Figure 5A highlights the FND120 (red) present within a central layer of the cell, by overlaying the FND z-stack across 3 μm with a single mid-plane optical slice of the cell stained for viability (blue). The elongated FND structures are an artifact from fluorescence bleeding through the z-layers.

**Figure 5.**
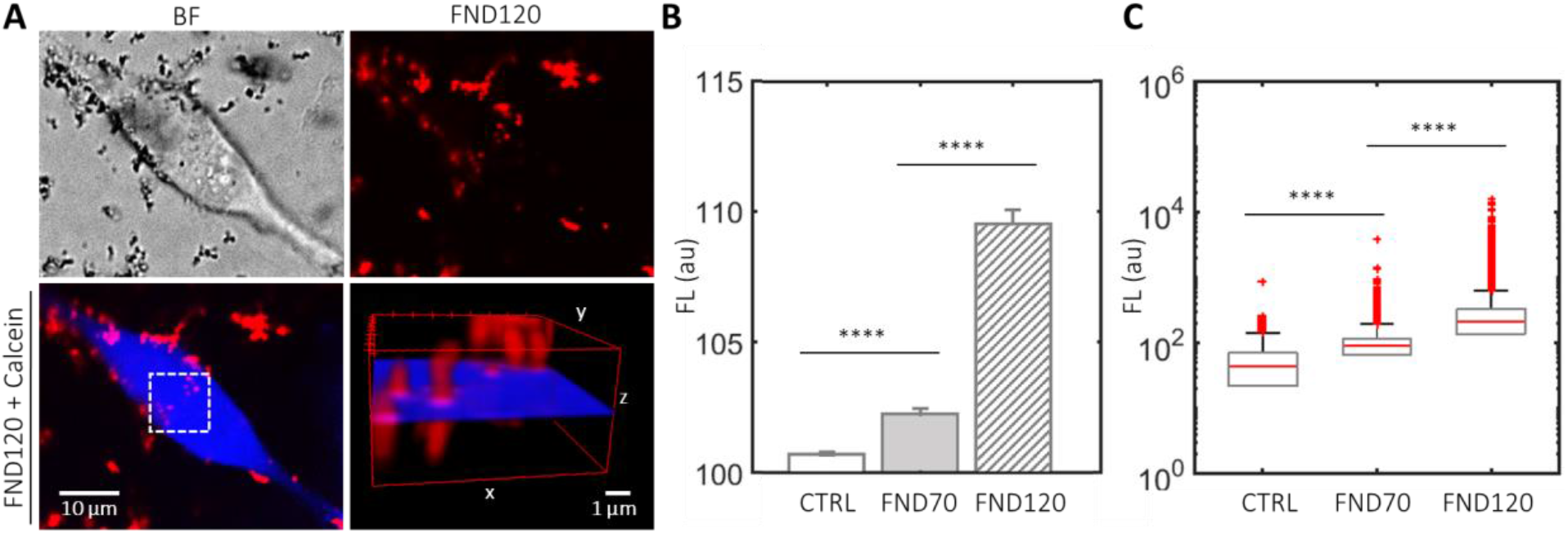
Cell uptake of fluorescent nanodiamonds (FNDs) by U87 cell monolayers, with FND diameters of 70 nm (FND70) and 120 nm (FND120). A) Mid-plane optical slices of a cell stained for viability (blue) and exposed to FND120 (red) for 30 min. Zoom in on cellular internalization. FND signal B) from cell monolayers, by image analysis (SD, n=10). Statistical significance: one-way ANOVA, and C) from dissociated cell monolayers for single cell analysis by flow cytometry (n= 500k). Statistical significance: Friedman’s test, a = 0.05, ****p < 0.0001.

Comparison of the cellular accumulation of FND70 vs FND120 was performed by quantitative image analysis 2 h after administration, where FND120 fluorescence signal appeared significantly stronger (p < 0.0001) (Fig.5B). The results are corroborated by single cell analysis using FACS on dissociated cell monolayers, confirming that FND120 uptake by U87 cells exceeded that of FND70 (Fig.5C, S3). Together, these results indicate increased accumulation and uptake of FND120 over FND70 in 2D U87 cell monolayers.

### Mass Density effect on differential NP – cell interactions in the 2D vs 3D microenvironmet

The discrepancy in size-dependency of FND uptake by U87 2D monolayers versus 3D spheroids is likely explained by the rapid FND sedimentation over the course of 2 h exposure, due to the large FND mass density. FND settling causes increasing FND effective-concentration in the 2D monolayers, and conversely decreasing FND effective-concentration in the 3D spheroids. It is important to note, that regarding the spatial resolution, the confocal microscope used for imaging cell monolayers surpasses the light-sheet microscope used for imaging spheroids. This suggests that, with the latter technique, larger FND accumulation was required for effective detection, thus it can be tuned by increasing either the FND effective-exposure-time or their effective-concentration. We herein, verified our results using a common method in both 2D and 3D models, i.e. FACS analysis, but we emphasize the importance of precise experimental documentation and careful data interpretation when comparing results across studies to progress translational research.

The tumor-in-a-tube platform involves an initial convective flow induced by NP administration that accelerates the early NP – spheroid interactions^40^, but thereafter the system transits into a more static state. In the absence of fluid dynamics, NP accumulation relies on passive diffusion for smaller particles, transcellular transport mechanisms and gravitational-force driven early-interaction. NP sedimentation proves critical in both 3D and 2D static cultures, but is reduced with fluid dynamics *in vivo* and on organ-on-a-chip microfluidic devices^59,60^. Though FND settling hinders their interaction with the 3D tumor spheroids, it enhances FND interactions with the 2D adherent cell monolayers.

Our study demonstrates that NP mass density is a key restrictive factor in NP-spheroid interaction dynamics. While NPs are often reported to interact differently with 2D adherent monolayers compared to suspension cultures, such as 3D tumor spheroids^61,62^, gravity is rarely acknowledged as the restrictive factor^63,64^. For NPs with large mass densities, gravity significantly alters the overall nanomaterial density (mass/ volume) in the course of time *in vitro*. We have previously discussed this gravitational effect with FND120 administration to a single-cell suspension monoculture, where similar to the 3D spheroids, the FND effective-concentration is increased within the compact cell arrangement in round-bottom well-plates compared to flat-bottom ones^3^.

## Conclusion

This study introduces a novel tumor-in-a-tube platform to monitor for the first time the dynamic interactions of fluorescent nanodiamonds (FNDs) with tumor spheroids. We employed 4D light-sheet microscopy and quantitative 3D radial analysis up to 100 μm towards the U87 spheroid core, and we addressed the challenges of dim FND fluorescence against cell autofluorescence. Uptake was validated by single-cell analysis on dissociated spheroids with FACS and in 2D cell monolayers.

We benchmarked FND accumulation against polystyrene nanoparticles (PNPs) at two nominal sizes each: 20 nm (PNP20), 70 nm (FND70), 100 nm (PNP100), and 120 nm (FND120). Overall, FNDs were detectable up to 50 μm spheroid depth, and they exhibited efficient accumulation, retention and a non-uniform, steep radial distribution profile. Conversely, PNPs achieved significantly enhanced uptake, retention, uniform-distribution and deeper spheroid convection, which however was size-dependent, in favor of the smaller PNP20. Unlike PNPs, FND120 outperformed the smaller FND70, although with low statistical significance. The discrepancy from size-dependent penetration is attributed to their fluorescence, which scales with particle size as larger FNDs contain more nitrogen-vacancy centers in their carbon lattice. Thus, the dim FND fluorescence necessitates greater FND agglomeration for smaller FNDs to reach detectable fluorescence levels. FNDs were also challenged with limited spheroid interaction due to rapid sedimentation, which minimized their effective-availability to roughly 30 min, 4-fold lower than the sustained effective-concentration by the lower-density PNPs. In 2D cell monolayers, gravitational FND settling promoted their cellular interactions resulting in size-dependent FND uptake, as confirmed through confocal microscopy and on dissociated monolayers using FACS. Moreover, FND treatment did not impair spheroid viability, while FND120 were localized beyond the plasma membrane within individual cells by 30 min exposure.

Altogether, our findings suggest FNDs as efficient, biocompatible long-term labels for U87 cell monolayers and spheroids, allowing for cell isolation from different spheroid layers. These properties make them promising candidates for biomedical applications such as real-time biosensing, long-term cell labeling and drug delivery in complex tumor environments.

Furthermore, our tumor-in-a-tube platform coupled with 4D imaging, provides a novel approach to understand the varying NP dynamics in 2D versus 3D environments. It highlights the importance of 3D models to better simulate *in vivo*, for NP assessment, and emphasizes the need of calibrating the NP effective-exposure-time in comparative NP assays, at 3D static models. These results underscore the need for careful experimental design and result interpretation, as well as detailed documentation of the experimental procedures. The application of the platform is relevant but not limited to nanoparticle applications, toxicology assessments, screening and time-pointing effective dosing, *in vitro*. Overall, our findings suggest that tailoring NP design based on mass density could optimize therapeutic delivery strategies and progress translational research.

## Materials and Methods

### Nanoparticles

The fluorescently-labeled anionic polystyrene nanoparticles (ThermoFisher, F8887; Exc/Em: 580/605 nm) were COOH-modified with nominal diameter sizes of 20nm and 100nm. The fluorescent nanodiamonds with negatively charged nitrogen-vacancy (NV) centers (Adamas Nanotechnologies, NDNV70nmHi, NDNV70nmHi; Exc/Em: 570/680 nm) were carboxyl-terminated with a negative surface charge and nominal diameters of 70nm and 120nm. FND120 had 3 ppm nitrogen-vacancies corresponding to approximately 400-500NV centers per particle, while FND70 had 2.5 ppm which corresponds to approximately 70-90 NV-centers per particle. All NDs were subjected to bath-sonication for 10 min before application.

The estimation of the travelling distance of a single NP undergoing sedimentation was based on a balance between gravity and friction from Stoke’s law:

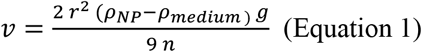

where *r* is the NP radius, *ρ*_*NPD*_ is the NP mass density, *ρ*_*medium*_ is the mass density of the medium, *g* is the gravitational acceleration and *n* is the viscosity of the medium.

The estimation of the total number of NV-centers in a single FND was based on the following formula:

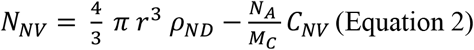

where *r* is the radius, *ρ*_*FND*_ is the FND mass density, *N*_*A*_ is Avogadro’s number, *M*_*C*_ is the molar mass of carbon and *C*_*NV*_ is concentration of NV centers.

### Cell Culture

Gliosblastoma multiforme U87-MG cell line (ATCC, HTB-14) was maintained in high glucose Dulbecco’s modified Eagle Medium (DMEM) supplemented with 10% Fetal Bovine Serum (FBS), 100 units/mL penicillin and 100 μgr/mL streptomycin (Gibco). Adherent cells were passaged upon 95% confluency, using TrypLE express (1X, Gibco).

### Gravitation-Assisted Spheroid Formation

Tumor spheroids were formed by gravitation-assisted method, seeding U87 cell suspension in culture medium (5,000 cells/mL) in round-bottom ultra-low attachment 96-well-plates (Corning Costar), at a volume of 100 μL/well. After 3 days of incubation at 37° C, 5% CO_2_, 100% humidity, multicellular tumor spheroids were harvested for experiments and were characterized with Matlab software (MATLAB R2020b; MathWorks). Spheroid viability was assessed by Fiji Software, where the positive control was treated with EtOH for 2 h (Figure S1).

### 3D Spheroids with FND Treatment for FACS assay

For single cell analysis, spheroids were treated with FND70 or FND120 (10 μgr/mL) for 2 h, and co-stained with cell-permeable nucleic acid stain (SYTO13, Invitrogen; Exc/Em: 488/509 nm) for 30 min, before harvesting. Spheroids were dissociated by trypsinization and agitation. The single-cell suspensions were pooled and fixed with 4% PFA. Cells were filtered through Falcon cell-strainers of 35 μm mesh (Corning), before examination with fluorescence-activated cell sorting (FACS, BD Biosciences; LSRFortessa). Cells Spheroids were additionally labeled with cell-permeable viability stain (calcein UltraBlue AM, Cayman Chemical; Exc/Em: 360/445nm) for FND biocompatibility assessment. We verified that cell fixation does not alter the cells in size (FSC) and granularity (SSC) (Figure S4). Single-cell gating, quadrant statistics and numerical analysis on the fluorescence values were executed with custom-scripts in Matlab software. For data loading we used the FCS data reader package^65^. When applicable, data is presented by the mean fluorescence intensity (MFI).

### 3D Spheroids for Dynamic Live-Cell Imaging by Light-Sheet Microscopy

Fully formed spheroids were rinsed 3 times with phosphate buffer saline (Gibco) and were supplemented with a cell impermeable dye (Alexa Fluor 488 Hydrazide, Invitrogen; Exc/Em: 493/517 nm). Each spheroid was positioned into a fluorinated-ethylene-propylene tube (inner diameter 0.8 mm, outer diameter 1.6 mm; Bohlender), which was then mounted on the microscope stage and submerged in heated water. Timelapse cell imaging was conducted with an Aurora airy light sheet microscope (M Squared, FOV 600 mm x 600 mm). The microscope is equipped with an M2 filter cube and two water-immersion objectives positioned perpendicular to one another, for illumination (UMPlanFL N 10x/0.30 W, WD 3.5 mm) and detection (UMPlanFL N 20x/0.50 W, WD 3.5 mm). To separately visualize the spheroids and the nanoparticles, laser beams for blue (488 nm) and yellow (570 nm) light were collected separately through a band-pass filter (520-540 nm) and long-pass filter (>570 nm), respectively. For each wavelength, we utilized the Aurora Alpha acquisition software to record a stack of 500 images (32-bit, 4 GB) with the same voxel size (0.2925 μm x 0.2925 μm x 0.4 μm) and imaging conditions (laser intensity/exposure time, minimized background light). For timelapse imaging of particle penetration into the tumor spheroids, a needle was inserted into the tube and positioned at a close proximity to the spheroid to administer the nanoparticles. An initial control set of stacks was recorded before particle introduction (t=-5 min). A second set of stacks was acquired immediately after nanoparticles were observed in the field of view (t=0 min), followed by another 24 sets of stack recordings with 5 min intervals, adding up to a dataset of 208 GB per spheroid. Temperature was constant at 37° C throughout the experiments.

### Image Processing and 4D Image Analysis of the Nanoparticle Penetration in Spheroids

We report a total of 17 spheroids, corresponding to 3.5 TB of data. Image processing requires a powerful computer. To apply 4D analysis across half a spheroid, we considered 100 μm (250 slices) from the surface layers towards the spheroid core in the axial direction (z-axis), for both wavelengths. The corresponding GFP stacks were subjected to image segmentation to localize the un-stained spheroid within the surrounding fluorescent buffer. This provided unique 3D binary images so that the subsequent FND intensity measurements are confined to the spheroid periphery. All holes were filled inside the masks. A radial analysis was performed on the RFP stack where the nanoparticles are visualized. Each radius calculates the line profile from the center of mass of the binary image towards the edge, and this was repeated over 180° and across the entire z-stack. We then limited the radii lengths to 100um from surface to center and the pixel values were averaged across all radii in the three dimensions. After obtaining the averaged 4D radial profile, we summed the values every 20 μm to get the 5 shells of the integrated densities. For each spheroid, the analysis was repeated on 26 timepoints, acquired during the time-lapse imaging.

Thereafter, we proceeded to data normalization. We first normalized the data by the signal from the control image of the spheroid before particle injection (t = -5 min), to compensate for uneven fluorescence due to the spheroid morphology. Next, to overcome the cell autofluorescence bleaching factor over time, we subtracted the 4D signal from control spheroids without FNDs, which exhibited a photobleaching decay. Normalization was particularly useful to decouple FND signal from the background noise. FNDs are dim particles whose fluorescence competes with the cell autofluorescence at low particle concentrations. While FND signal increased with time, cell autofluorescence decreased by photobleaching, which are two competing processes.

### 2D monolayers with FND Treatment for Live-Cell Imaging and FACS assay

For both assays, cells were seeded for 24 h (25,000 cells/cm^2^), in 35 mm glass-bottom Petri dishes at 37° C, 5% CO2, 100% humidity, to achieve a 70-90% confluency before FND application (10 µgr/mL). Microscopy imaging was conducted using a spinning-disk (SD) confocal-microscope (Nikon Ti2) with an oil immersion objective (CFI Plan Apochromat Lambda, 60x/1.4, oil, WD 0.13 mm). Laser beams for near-UV (405 nm) and yellow (561 nm) light were collected used for excitation of the viability stain and FNDs, respectively. Images (16-bit) had voxel size (0.1828 μm x 0.1828 μm x 0.3 μm) and were analyzed using Fiji software. For FACS, cells were dissociated with trypsin and fixed with 4% PFA.

### Statistical Analysis

Statistical analysis was performed with Matlab software. Statistical significance was evaluated by one-way analysis of variance ANOVA or Friedman’s test, as stated under each figure, at level of significance α = 0.05, with p-values *p < 0.05, **p < 0.01, ***p < 0.001- and ****p < 0.0001.

## Acknowledgements

This work was supported by Independent Research Fund Denmark (grant no 0135-00142B) and the Novo Nordisk Foundation (grant no NNF20OC0061673).

## Author Contributions

MN designed the study under supervision of LJ, MD, and KBS. MN conducted the experiments, analyzed the data and drafted the manuscript. All authors revised and approved the manuscript.

